# The Microbiota’s Response to Host Adaptive Evolution

**DOI:** 10.1101/2025.09.22.677868

**Authors:** Zachary S Greenspan, Robert Courville, Courtney Mueller, Kenneth R Arnold, Kathy Thien, Michaelangelo Marcellana, Han Yin, Arnie Lynn C. Bengo, Kshama E Rai, Elnaz Bagheri, Sam Behseta, John Chaston, Parvin Shahrestani

**Author notes:** Corresponding author: Parvin Shahrestani, 800 N State College Blvd, Fullerton CA 92835.

## Abstract

Over the past two decades, experimental evolution has significantly advanced our understanding of evolutionary patterns and mechanisms of genomic change. One critically underexplored dimension is the role of microbiota in host adaptive evolution. In this research we investigate the microbiota from forty laboratory-selected *Drosophila melanogaster* populations exhibiting four distinct aging trajectories, ranging from extremely-short lifespans to -long lifespans. Using metagenomic sequencing and colony forming unit (CFU) counts in both conventional and gnotobiotic conditions, we uncover substantial microbiota differentiation among these populations. The most striking pattern is the consistent loss of *Wolbachia* in populations selected for rapid aging, in contrast to its near complete dominance in long-lived populations. This suggests a positive association between relative abundance of *Wolbachia* and lifespan, alongside a negative correlation between *Wolbachia* and Acetic Acid Bacteria (AAB) titres. These findings position the microbiota, and particularly *Wolbachia*, as a potentially integral component in host life history evolution, with implications for understanding the microbial contributions to aging and adaptation.

**Significance:** Fruit flies bred for short lifespans consistently lose the bacterial symbiote *Wolbachia*, while the microbiota of long-lived flies are almost entirely dominated by it. There is a stable, recurring link between this microbe and lifespan, and traits associated with lifespan. This persistent pattern suggests that microbiota, *Wolbachia* in particular, may play a direct role in shaping the evolution of lifespan.

## Introduction

The interaction between environmental and genetic factors in shaping an organism’s traits is complex and multifaceted. This complexity can be explored in model systems such as the fruit fly *Drosophila melanogaster*, which harbor substantial phenotypic and genetic variation for life history traits, making these traits particularly responsive to natural selection. For example, laboratory selection on age-of-first-reproduction in populations of *D. melanogaster* quickly leads to divergence, not only in genome-wide patterns but also in traits such as developmental rate, aging, lifespan, fecundity, and stress resistance (Rose et al., 2004). One critical but underexplored aspect of *D. melanogaster* laboratory selection studies is the role of microbiota, the microorganisms residing on or within the host. In *D. melanogaster*, microbiota can substantially influence host traits, including life history traits such as development and aging (Matthews et al., 2020; Walters et al., 2020b), and shape the flies’ life history evolution (Rudman et al., 2019). Nevertheless, it is unknown how laboratory selection on life history traits impacts microbiota in *D. melanogaster*.

The Drosophila Experimentally Evolved Populations (DEEP) Resource, also known as the Rose populations, is a collection of populations that have evolved phenotypic and genomic divergence as a result of laboratory selection on life history traits (Rose et al., 2004). These populations have been maintained at large population sizes under consistent dietary conditions for hundreds of generations and exhibit repeatable patterns of divergence for life history traits. Populations selected for early reproduction show accelerated development and reduced lifespan compared to populations selected for late reproduction (Burke et al., 2010). These divergence patterns resemble clinal variation in wild populations of *D. melanogaster, with populations* from higher latitudes exhibiting larger body size, delayed reproduction, and greater stress resistance, compared to those from lower latitudes. Clinal variation in *D. melanogaster* populations has been associated with divergence of their microbiota (Walters et al., 2020a). However, the degree to which their microbiota differences are environmentally driven or genetically mediated remains an open question. The DEEP Resource offers a robust system for investigating the impact of host adaptive evolution on the microbiota, which provides an opportunity to dissect the role of host-microbiota interactions in shaping life history traits.

The *D. melanogaster* microbiota is of low numerical and taxonomic complexity, comprising approximately 20-100 distinct microorganisms that are primarily lactic acid bacteria (LAB) acetic acid bacteria (AAB), and Enterobacteriaceae {Ludington & Ja, 2020; Yu & Iatsenko, 2020; Staubach et al, 2013; Wong et al., 2011; Wong et al., 2013; Adair et al., 2018; Chandler et al., 2011}. The endosymbiont *Wolbachia* is also common in *Drosophila* lines, including laboratory lines (Clark et al., 2025; Bourtzis et al., 1996}. The *D. melanogaster* microbiota can be shaped by both environmental factors and genetic influences {Gale et al., 2024; Chaston et al., 2016} and can be transmitted to larvae hatching from eggs via the diet (Broderick et al., 2014; Blum et al., 2013}. Laboratory populations of *D. melanogaster* tend to bear microbial communities that have fewer bacterial taxa than wild populations {Staubach, 2013}. While most of the *D. melanogaster* microbiota can be cultured in the lab, intracellular *Wolbachia* cannot. Nevertheless, *Wolbachia* have been shown to influence host fitness, reproduction, and development (Bourtzis et al., 1996; Yu & Iatsenko, 2020).

We evaluated microbiota abundance and composition in experimentally-evolved *D. melanogaster* populations that have diverged for life history and stress resistance traits. Laboratory selection on host traits led to substantial repeatable changes in microbial communities, most notably, a consistent loss of *Wolbachia* in populations selected for early reproduction. Our findings provide evidence for host genetic control over the microbiota composition and abundance and suggest that this trait itself may evolve under selection.

## Methods

We analyzed microbiota divergence among experimentally evolved populations of *D. melanogaster* (Figure 1) to investigate how host genetic background and evolutionary history influence microbiota composition and abundance. We divide our analyses into two main experimental categories: Experiment 1, Sequencing, and Experiment 2, Direct Characterization.

**Figure 1.**
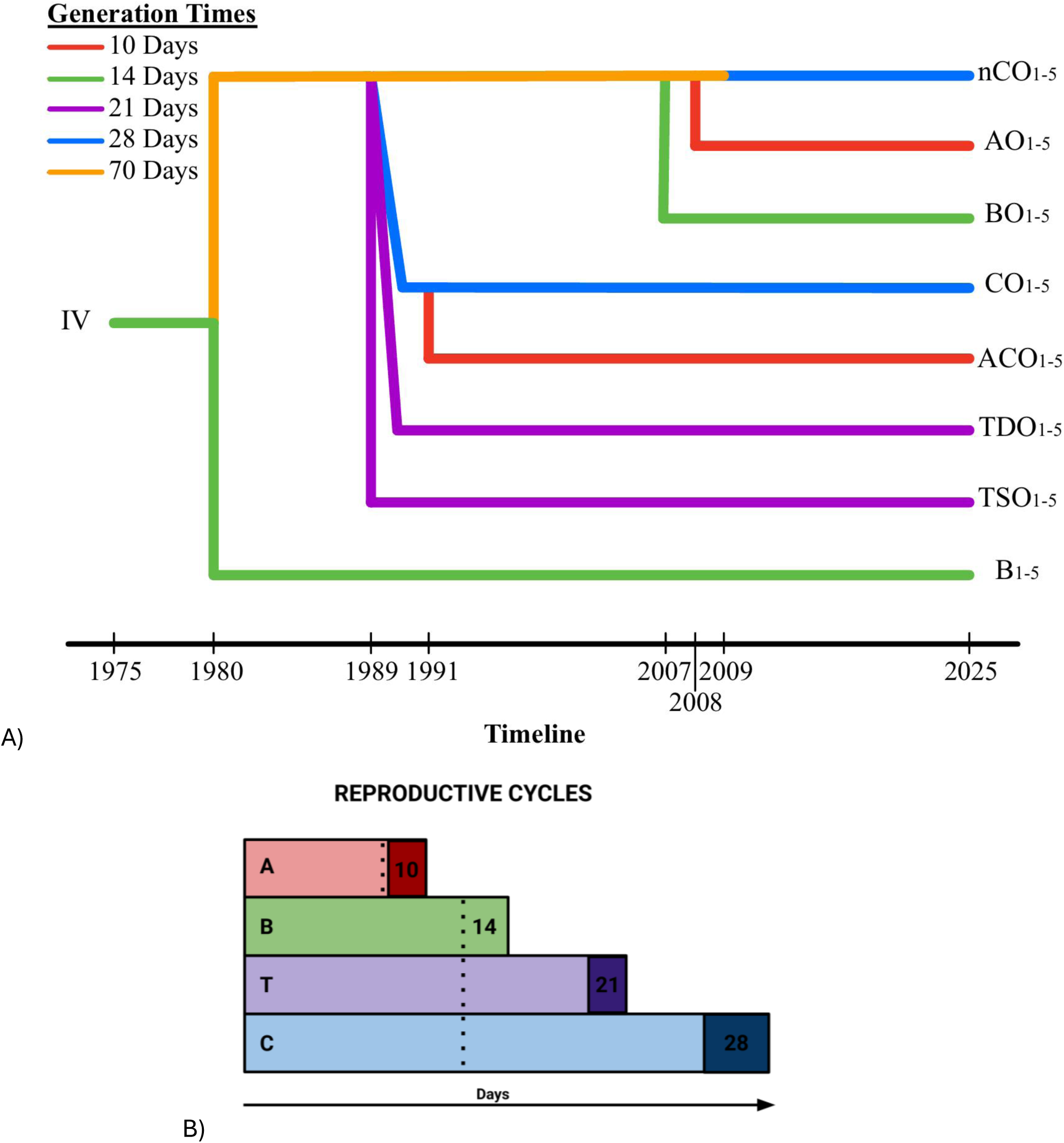
A) Phylogeny of *D. melanogaster* Populations Selected for Different Age of First Reproduction. The original IV population was collected back in 1975. O_1-5_ (shown in orange but are no longer maintained in this laboratory system) and B_1-5_ were derived from this initial IV population. The O_1-5_ populations gave rise to the TDO_1-5_, TSO_1-5_, CO_1-5_, AO_1-5_, BO_1-5_, and nCO_1-5_ populations. ACO_1-5_ populations were derived from CO_1-5_. Each color line represents the respective generation time each population is selected for. A change in color of the line indicates that a population underwent a change in its selection regime. The number of generations that populations underwent their most recent selection regimes are summarized in Table S1. Across the bottom are the dates that mark the start of any new selection regimes for any population. Subscripts 1-5 indicate that each population cohort is five-fold replicated. B) Graphical representation for the reproductive life cycle of each population type as defined by their selection for unique reproductive windows. Each dotted vertical line represents the transition from vials to cage while the darker sections represent the period of time flies are given yeast to facilitate egg laying before their selected date of reproduction.

### *Drosophila melanogaster* Populations

A wild *D. melanogaster* population with substantial additive genetic variation in life history traits was originally collected from an apple orchard in Massachusetts, moved to the lab and maintained on 14-day discrete generation cycles as the “IV” population (Ives, 1970; Rose and Charlesworth, 1980). Multiple populations of *D. melanogaster* descended from the IV population, forming the Drosophila Experimental Evolved Populations (DEEP) Resource (Figure 1). The DEEP resource is maintained on a banana-molasses diet (per 1L of distilled H_2_O: 13.5 g Apex Drosophila agar type II, 121 g peeled ripe banana, 10.8 mL Light Karo corn syrup, 10.8 mL Dark Karo corn syrup, 16.1 mL Eden organic barley malt syrup, 32.3 g Red Star active dry yeast, 2.1 g Sigma-Aldrich Methyl-4-hydroxybenzoate (an anti-fungal agent), and 42.5 mL EtOH), which has been consistent throughout their selection protocol and recent evolutionary history. The macronutrient composition per 1L of the banana-molasses diet is as follows: total fat = 1.2 g, sugar = 37.2 g, total carbohydrates = 90.36 g, protein = 21.2 g, and calories = 450 Kcal (Rutledge, 2018).

Both the O_1-5_ and B_1-5_ populations (with subscripts indicating the five replicates) were derived from the original IV population. The O_1-5_ populations were selected for late-age reproduction with a 70-day generation cycle (Rose, 1984), while the B_1-5_ populations were selected for early-age reproduction with a 14-day generation cycle (same as the IV). In 1988, the O_1-5_ populations gave rise to the TDO_1-5_ and TSO_1-5_ populations (Rose et al., 1992). The TDO populations were intensely selected for desiccation resistance for ∼260 generations. Concurrently, the TSO populations served as controls for the TDO populations, undergoing moderate selection for starvation resistance over the same period. The TDO and TSO populations then underwent ∼240 generations of being maintained on a relaxed 21-day selection before sequencing (Phillips et al., 2019). In 1992, the O_1-5_ populations were used to create the CO_1-5_ populations, maintained on a 28-day cycle (Chippindale, 1996). These CO_1-5_ populations were then used to create the ACO_1-5_ populations, whose generation cycles were shortened to 10 days in 1997 (Chippindale, 1996).

In the mid 2000s (Figure 1), the AO_1-5_ (with a 10-day generation cycle), BO_1-5_ (with a 14-day generation cycle), and nCO_1-5_ (with a 28-day generation cycle) populations were rederived from the O_1-5_ populations.

ACO_1-5_ and AO_1-5_ will henceforth be called A-types, B_1-5_ and BO_1-5_ will henceforth be called B-types, TSO_1-5_ and TDO_1-5_ will henceforth be called T-types, and CO_1-5_ and nCO_1-5_ will henceforth be called C-types.

Before initiating any assays, all populations were reared under identical conditions under a continuous 24-hour light exposure at a constant room temperature of 25°C and at controlled densities (approximately 60-80 larvae per vial) for two 14-day generations to mitigate environmental and parental effects. Due to differences in development time between population types, these lead-in generations were staggered to enable time-matched bacterial quantification of 5-7-day-old flies post eclosion.

### Metagenomics Analysis of Aging Populations from New and Extant Genomic Datasets

Some of the whole genome sequence data analyzed herein was obtained from prior publications on these populations (Graves et al. 2017 for the A-, B-, and C-type populations, and Phillips et al. 2018 for the T-type populations). In these prior works, DNA was extracted from pooled samples of 120-200 whole-body homogenized female flies using the Qiagen/Gentra Puregene kit as per the manufacturer’s protocol for bulk DNA purification. The genomic DNA pools were prepared as standard 200-300 bp fragment libraries for Illumina sequencing, with each set of five replicate populations of a treatment (e.g., ACO_1-5_) given unique barcodes, normalized, and pooled together. Each population was sequenced twice, and data from both runs were combined for our analysis.

In November 2018, samples of 200 female flies were collected for genomic sequencing from each of the 20 individual populations (ACO_1-5_, AO_1-5_, CO_1-5_, nCO_1-5_) and extracted using Qiagen Puregene kits. Each cage-cohort of ∼1,500 flies was generated from each of the five replicate populations for ACO, AO, CO, and nCO and maintained on a 14-day culture cycle for two lead-in generations, staggered a day apart to mitigate the impact of gene-by-environment interactions. The female flies were immediately dry frozen in liquid nitrogen and stored at −80°C and homogenized in liquid nitrogen prior to extraction. The 20g DNA pools were prepared as standard 200-300 bp fragment libraries for Illumina sequencing and constructed such that each five replicate populations of a treatment (e.g., ACO_1–5_) were given unique barcodes, normalized, and pooled together. Each 5-plex library was run on individual PE150 lanes of an Illumina NovaSeq6000 S4 at the UC Irvine Genomics High-Throughput Facility (GHTF). Resulting data were ∼100 bp paired-end reads. One specific population from this dataset, ACO1 was additionally sequenced at higher depth than (resulting in ∼40x reads processed for later metagenomic analysis).

The 16S rRNA marker analysis followed an established protocol (Kozich et al., 2013) and was performed as in previous studies (Rudman et al., 2019; Mabey et al., 2020; Walters et al., 2020b). Briefly, DNA was extracted from pools of 20 whole body flies using the Zymo Research Quick-DNA Fecal/Soil Microbe 96 Kit (D6010; Irvine, CA). The V4 region of the 16S rRNA was PCR-amplified using dual-barcoded primers (Kozich et al., 2013) and Accuprime PFX polymerase (ThermoFisher Scientific, Waltham, MA, USA). Amplicons were normalized and purified, and sequencing was performed using v2 chemistry on a partial lane of a HiSeq 2500 (Illumina, Inc., San Diego, CA) at the BYU DNA sequencing center.

### Phenotypic Analysis through Gut Characterization and Bacterial Load Analysis

All phenotypic analysis was done using ACO_1-5_ and CO_1-5_. For each individual population replicate, 5-7 days following eclosion from pupa casings, pools of five female and five male flies were sampled from their vials and placed into a microcentrifuge tube containing 125 μL of mMRS (modified De Man, Rogosa, Sharpe agar) medium and ∼0.125 g of MP Biomedicals® Lysing Matrix D Bulk beads (Rudman et al., 2019). The pools of flies were then homogenized at a speed of 4.0 m/s for 30 seconds each using a MP FastPrep-24 homogenizer, before 875 μL of mMRS was added to each homogenate. For each population replicate, 10 sets of five flies from each sex for a total of 100 flies were homogenized. Serial dilutions of 1:1, 1:8, 1:64, 1:512 of the homogenates were made using mMRS broth and plated on mMRS agar plates in streaks separated by dilution. Homogenates from each set of five flies were plated on two mMRS agar plates: one for aerobic growth of AAB, and one for the anaerobic growth of LAB. Agar plates for anaerobic growth were placed into an airtight container and flooded with CO_2_ to rid the container of any oxygen. Each treatment condition was then kept inverted (lid down) for 48 hours at 30°C for colony growth. Colonies were then analyzed by morphology and classified as either AAB or LAB, following protocols of Koyle et al., 2016. 20 flies per sex per population replicate were individually weighed, and those weights were averaged. The colony forming units were then divided by the average weights for each population replicate to generate a CFU fly mg^-1^ count.

Gnotobiotic flies were reared by inoculating sterile eggs with defined microbial communities as described previously {Koyle, 2016}. Briefly, the bacterial inoculum was prepared by culturing two AAB strains - *Acetobacter pomorum* DmCS_004, *A. tropicalis* DmCS_006 - and two LAB strains - *Levliactobacillus brevis* DmCS_003, *Lactiplantibacillus plantarum* DmCS_001 - in pure culture on mMRS medium at 30°C for 48 hours. LABwere incubated in a microoxic environment created by flooding an airtight container CO_2_ Then, the bacterial cultures were normalized to OD_600_ = 0.1. For normalization, 200μL of each bacterial culture was transferred to a 96-well plate to create serial dilutions, and the OD_600_ of each dilution was measured using a Thermo Scientific™ NanoDrop™ One Microvolume UV-Vis Spectrophotometer. The bacteria were centrifuged and resuspended in PBS at OD_600_ = 0.1.

*D. melanogaster* embryo collection plates were inserted into each population cage approximately 18 hours before dechorionation. In a Laminar flow hood, the eggs were rinsed into a sieve lined with fine mesh using distilled water, and then rinsed three times in a 0.6% sodium hypochlorite solution for 2 minutes and 30 seconds, followed by three separate dips in autoclaved distilled water for 30 seconds each. Successful dechorionation was confirmed by the embryos adhering to the bushing walls. Dechorionated embryos were transferred at a density of 30-60 eggs per vial into a 50mL conical tube with sterile banana diet and lids slightly loose, and 50 mL of the prepared bacterial inoculant was added to each tube. For monoassociations, individual strains were inoculated to the sterile eggs; for gnotobiotic 4-species mix, equal volumes of normalized bacteria were pooled together, then inoculated to the sterile eggs.

To assess differences in gut sizes between the five ACO and five CO populations, we dissected the guts of adult flies (Bourtzis etal., 2000) from both sexes and compared the measurements to body and abdomen measurements. Flies were anesthetized using a Carolina Biologics FlyNap anesthetic kit as per the recommended protocol, by exposing them to a wand soaked in the solution placed in the rearing vials for each population. Once anesthetized, male and female flies were separated, and measurements of body size and abdomen length were taken. Subsequently, the head of the fly was removed from the body, and the gut was carefully extracted through the bottom of the abdomen, ensuring that the crop and Malpighian tubules were identified to verify complete gut extraction (Broderick et al., 2014; Miccheli et al., 2015). Each experimental replicate involved 80 dissections and measurements per replicate, with 20 per sex per population. The experiments were conducted in triplicate, resulting in a total of 240 dissections per treatment replicate. Two treatment replicates were performed, totaling 480 dissections and measurements. Ratios of gut to body and gut to abdomen were then calculated for each sex and population.

### Metagenomics Analysis of Aging Populations from New and Extant Genomic Datasets

From previous experiments, we have Illuminia short read sequence data from 10 A-type populations (Graves et al., 2017), 10 B-type populations (Graves et al., 2017), 10 T-type populations (Phillips et al., 2018), and 10 C-type populations (Graves et al., 2017). These populations encompass an aging-genotype gradient dataset.

Additionally, the 10 A- and C-type populations were resequenced in 2018. Compared to the genome data from Graves et al. 2017, the 2018 sequence data occurred after 250 and 90 further generations for the A- and C-types respectively, allowing us to compare the relative abundances of bacteria for the same populations across chronological time and generation count. This comparison encompasses a time-displacement population dataset.

Read matching and analysis of all genomic data collected were performed using MetaPhlAn 4.0.6 with the following parameters “--index mpa_vJun23_CHOCOPhlAnSGB_202403 --nproc 20 --stat_q 0.”, which matched the data with known bacterial genome sequences (Blanco-Míguez 2024).

We used genus as our phylogenetic grouping of choice due to the loss in fidelity at the species level where it became too difficult to make comparisons between populations. Ie: while 35 samples feature Weisella, 2 samples feature *Weissella cibaria*, 2 samples feature *Weissella hellenica*, 10 samples feature *Weissella minor*, and 35 samples feature *Weissella viridescens*. Subsequently, it makes more sense to go one level higher to not dilute the detection of any potential effects on abundance levels if we had analyzed at a classification level where proper comparisons could not be made.

For all analyses involving individual genera, we filtered out any genus present in our dataset with less than 1% overall presence across all samples. This identified four genera: *Wolbachia* (48.14%), *Weissella* (22.97%), *Acetobacter* (21.86%), *Lactiplantibacillus* (6.261%), which account for over 99.2% of all metagenomic bacteria present in our samples.

Because such methods only allow for detection of overall categorical differentiation rather than pairwise differentiation, some common methods such as linear models and Anovas lack utility in answering our hypotheses. Therefore, we used pairwise Wilcoxon Rank Sum tests as implemented by the rstatix R package (Kassambara 2023) for individual bacteria by population level comparisons. For Wilcoxon Rank Sum tests, we used Benjamini Hochberg (1995) false discovery rate correction to account for multiple comparison testing when evaluating significance between pairs. We ran 1000 permutations testing for significance between pairwise comparisons and results were internally consistent. Out of the 1000 permutations and related pairwise comparisons (6000 tests total), 5 were as or more significant than our results presented.

We used the pheatmap package in R (Kolde 2022) to generate a heatmap with clustering from treating each population independently for similarity in the patterns of bacterial relative abundances.

Lastly from metagenomics, we compared the MetaPhlan results from 27,710,275 (27 million) ACO_1_ DNA reads to 1,045,075,688 (1 billion) ACO_1_ DNA reads within the 2018 *Drosophila* sample to see if we could detect the presence of *Wolbachia* at extremely deep sequencing depths.

Data analysis on 16s rRNA markers were conducted using QIIME2 and R. Reads were trimmed, denoised, dereplicated, and called amplicon sequence variants using DADA2, and taxonomy was assigned with the GreenGenes classifier 13_8_99 (McDonald et al., 2012). ASV tables were filtered, and differences between ACO and CO treatments were determined by PERMANOVA (Oksanen et al., 2017) of Bray Curtis, weighted Unifrac, or unweighted Unifrac distances (Lozupone et al., 2005).

### Phenotypic Analysis through Gut Characterization and Bacterial Load Analysis

We used linear mixed effect models to evaluate the effects of sex and selection regime on CFU per fly abundances. All LMM analyses conducted in the paper used the lme4 package (Bates et al. 2015) in R (R Core Team, 2023). For each sample we have assayed the number of colony forming units *C*_*ijk*_, from selection regime (*i=*1 [ACO], 2 [CO]), sex (*j*=1 [Male], 2 [Female]), and population (*k*=1,2,…10). Values of CFU per fly are either in raw values or normalized by weight. By combining the abundance levels of both types of CFUs (Acetic Acid Bacteria and Lactic Acid Bacteria) into one dataset, we will also employ a microbe clade variable (*m*=1 [Acetic Acid], 2 [Lactic Acid]) in our linear model.

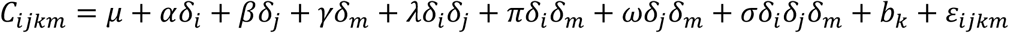

The random components, *b_k_* and ε*_ijk_* correspond to the random variation from population, and residual variation, respectively.

Or more straightforwardly:

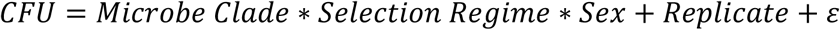

The methods for analysis of Gnotobiotic Flies were the same as the ones for CFU with one minor difference. Here the random effect is nesting replicate into experiment rather than simply replicate by itself given this specific experiment and assay was repeated 4 times.

We used linear mixed effect models to evaluate the effects of sex and selection regime on gut sizing. For each sample we have assayed the number of colony forming units *G*_*ijk*_, from selection regime (*h=*1 [ACO], 2 [CO]), sex (*i*=1 [Male], 2 [Female]), population (*j*=1,2,…6), sample (*k*=1,2,…*n*), and researcher (*m*=1,2). Values of CFUs are either in raw values or normalized by weight.

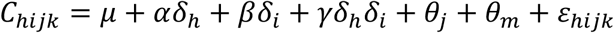

θ_*j*_ and θ_*m*_ represent the random effects by replicate and researcher respectively. Or more straightforwardly:

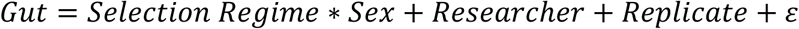

## Results

### Metagenomics Analysis of Aging Populations

In total, we analyzed microbiota data from 40 *D. melanogaster* populations across four selection regimes (A-, B-, C-, and T-types), with 20 of these populations (A- and C-types) resequenced in 2018 in addition to published datasets from prior studies. From our 60 pFour bacterial genera—Wolbachia, Weissella, Acetobacter, and Lactiplantibacillus—accounted for 99.2% of all bacterial reads, and analyses were conducted at the genus level to avoid species-level classification loss.

From these reads, we first investigated the non-reproductive tract microbiota of the different populations by filtering out reads assigned *Wolbachia* (Figure 2B). From this we compared the differentiation between selection regime on the non-*Wolbachia* genera specified above. First, *Lactiplantibacillus* shows no differentiation between any population-level comparison (Supplemental Figure 2). Acetobacter and *Weissella* relative abundances were the same in the B-, and C-type populations. However, the A-type populations had higher relative abundances of Weissella and lower relative abundances of Acetobacter compared to the B-, T-, and C-types. In contrast, T-type populations had lower relative abundances of Weissela and higher relative abundances of Acetobacter compared to the A-, B-, and C-types (Supplemental Figure 2).

**Figure 2:**
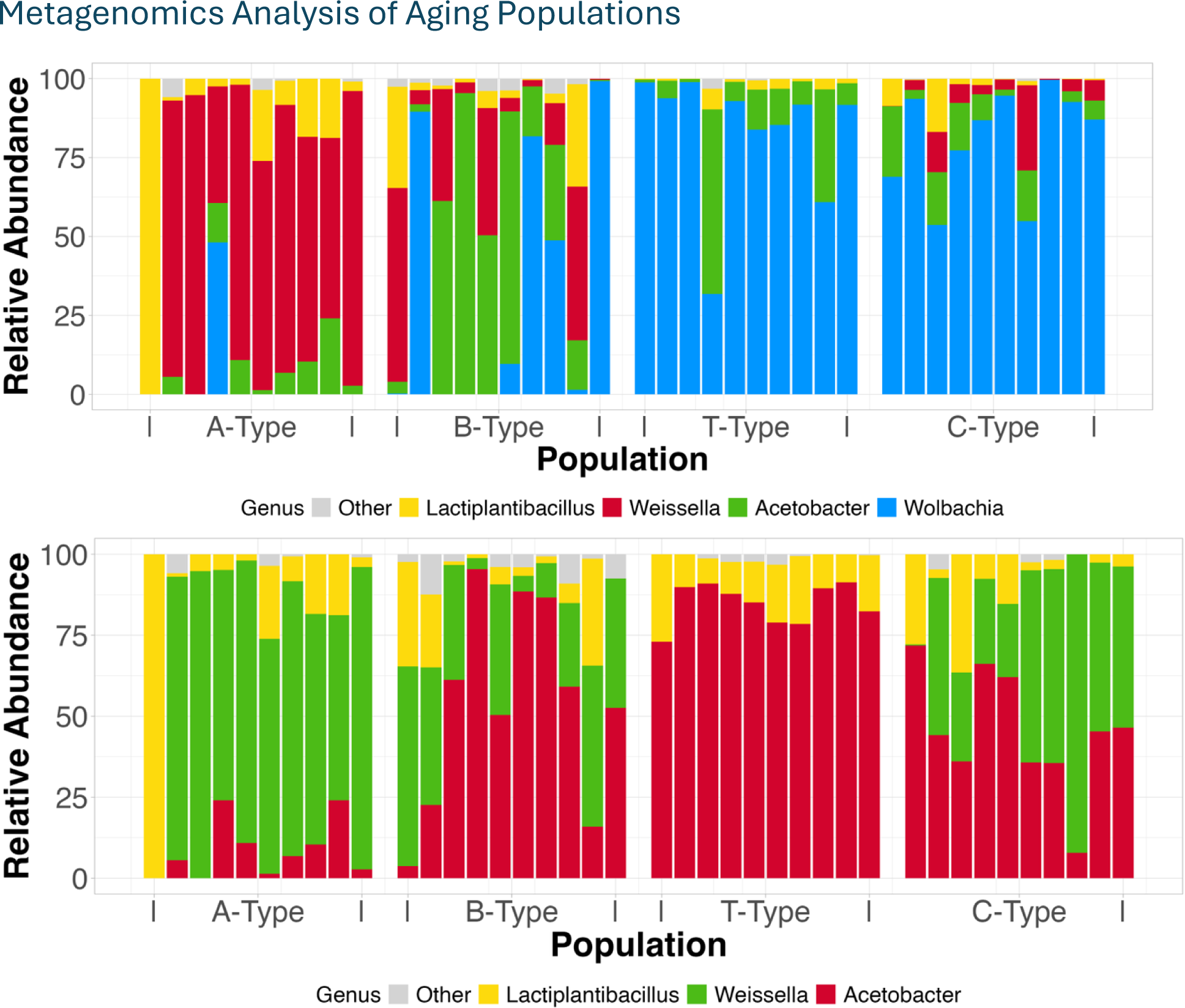
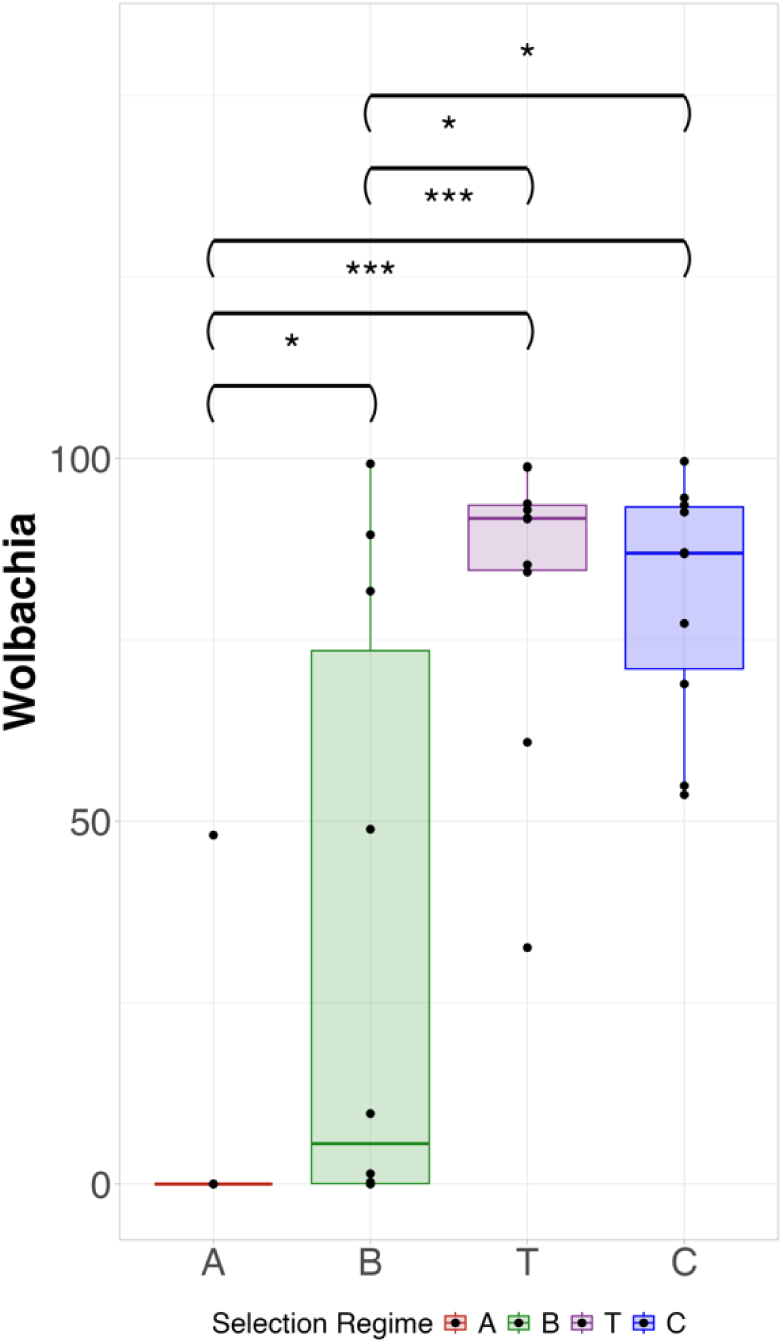
Relative abundance of bacterial clades from metagenomic data on the set aging-gradient populations from shortest lived (left) to longest lived (right). A) The abundance for all relevant genera. B) The relative abundance for any other genera after Wolbachia has been factored out. C) Wilcoxon rank-sum test for bacterial metagenomics abundances from metagenomic data on the set of aging-gradient populations from shortest lived (left) to longest lived (right) for the relative abundance of Wolbachia.

Turning to the relative abundance of *Wolbachia,* shorter-lived populations feature less *Wolbachia* than longer-lived populations (Figure 2A). Hierarchical clustering of samples according to their microbiota composition generally recapitulated the evolutionary history of the lines, further supporting the stark, *Wolbachia-*driven differences between the lines (Supplemental Figure 3). These trends were confirmed by resequencing additional collections of A- and C-type lines via each of whole genomic shotgun analysis and 16s rRNA V4 analysis (Fig 1C, Sup. Figure 1). Remarkably, the differences between A- and C-types remain stable: A-type populations consistently show minimal or no Wolbachia presence, while C-types how a substantial Wolbachia abundance (Figure 3A, 3C). This long-term consistency supports the hypothesis that *Wolbachia* abundance is a stable and defining feature shaped by selection regime. To test if the complete absence of Wolbachia detected in the A-types was merely a consequence from too low a read count, we investigated a single 2018 population sample at extremely deep sequencing. Even with deep sequencing at almost forty times a sequencing depth as the original sample, we were still unable to detect any presence of Wolbachia for an A-type population in 2018. Overall, our results provide evidence that specific selection regimes are associated with distinct microbiome profiles.

**Figure 3:**
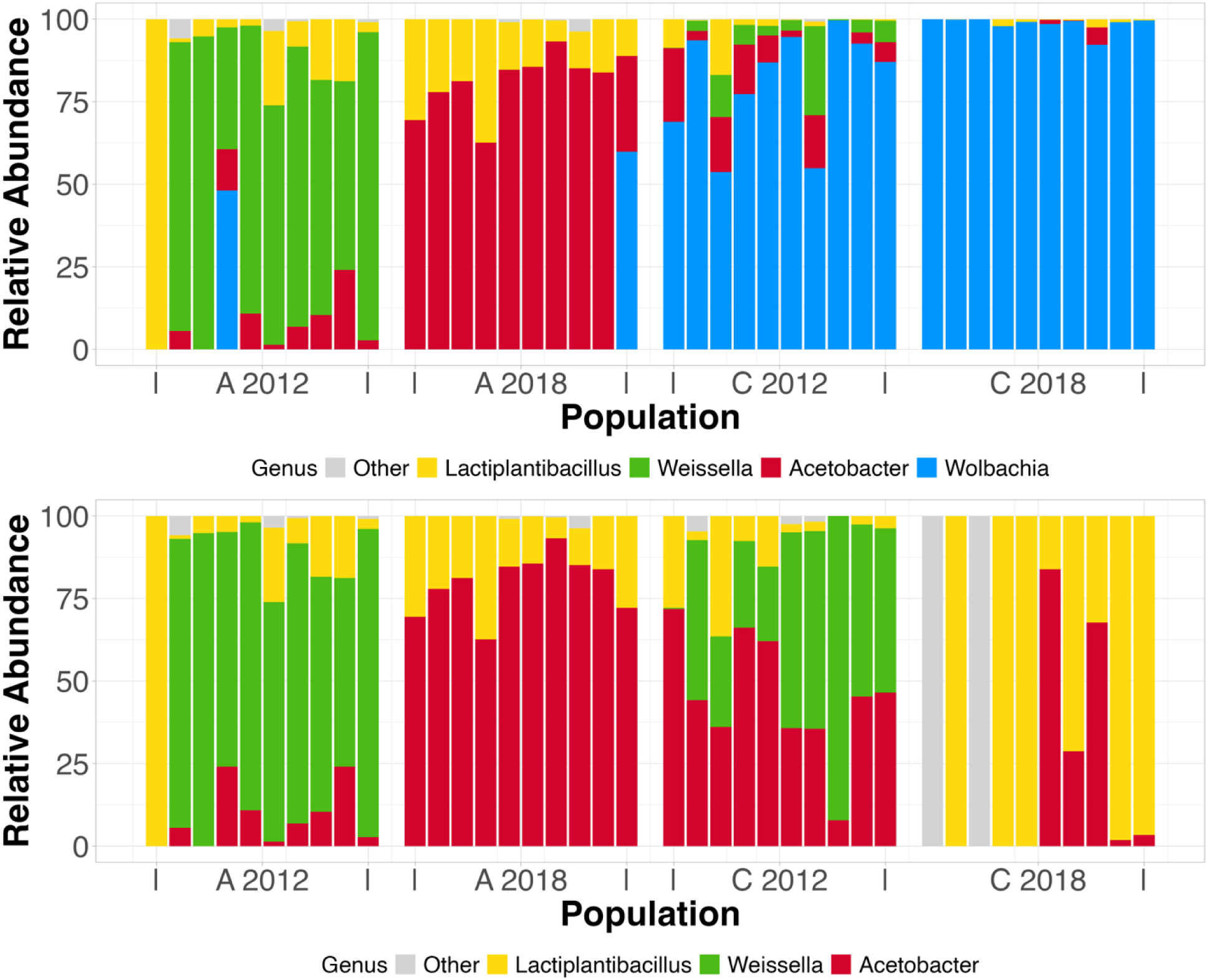
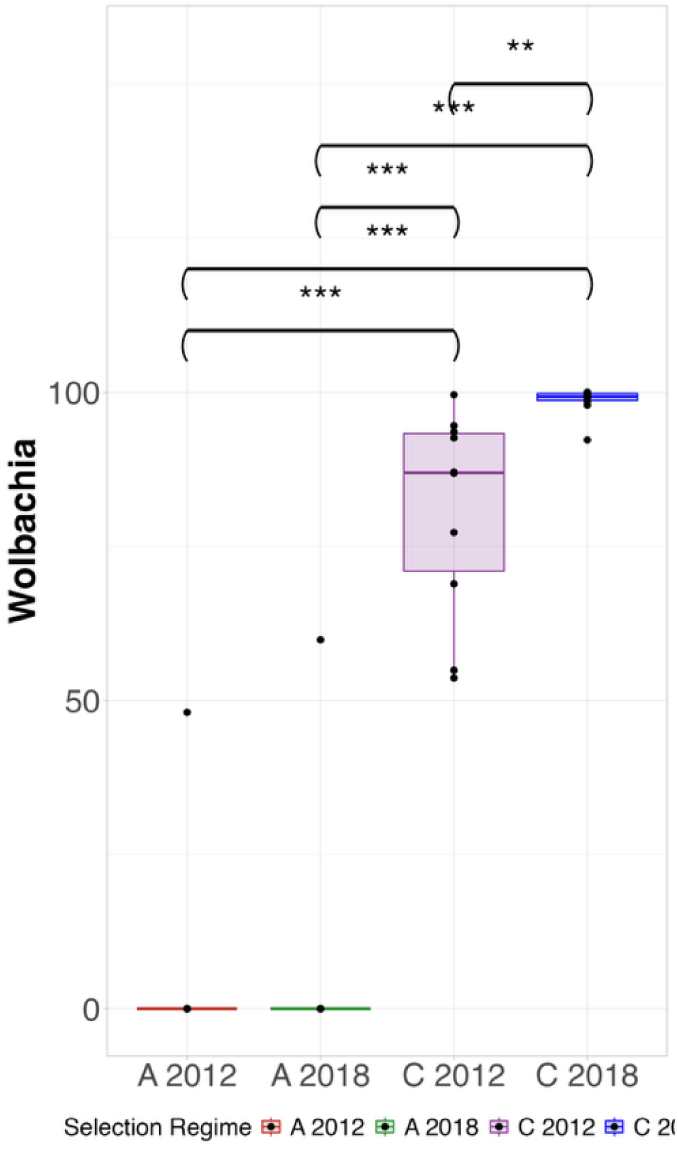
Relative abundance of bacterial clades from MetaPhlAn metagenomics on time and generation displaced A-type and C-type populations. There were 250 generations between A-type time points samples and 90 generations between C-type time points samples. A) shows the abundance for all relevant genera. B) shows the relative abundance for the other genera after Wolbachia has been factored out. Note: CO1 and CO3 genomics collected in 2018 registered as 100% Wolbachia which is why there are no genera represented once Wolbachia has been factored out. C) Wilcoxon rank-sum test for bacterial metagenomics abundances from metagenomic data on the time-displaced populations for the relative abundance of Wolbachia.

### Phenotypic Analysis through Gut Characterization and Bacterial Load Analysis

Metagenomic analyses allow us to detect the presence and relative abundance of bacteria, but do not reveal absolute bacterial loads or phenotypic consequences. To address this, we conducted direct phenotypic characterizations using colony-forming unit (CFU) counts to quantify bacterial abundances in our populations.

Across all statistical tests, we observed significant differences in CFU per fly between LAB and AAB, as well as between ACO and CO selection regimes (Table 1). These differences were consistent in both raw CFU counts and CFU counts normalized by fly weight (Sup. Tables 2). We investigated the CFU per fly of LAB and AAB independently, both with their raw amounts (Figure 4) as well as amounts normalized by fly weight (Supplemental Figure 4). When analyzing LAB and AAB CFU data independently from each other, we find the effect of selection regime to be significant in all cases, albeit less significant in AAB. We don’t find any effect of sex or the interaction of sex and selection regime on CFU per fly (except for a minor effect of sex on raw counts of LAB).

**Table 1:** P-values from a linear model using CFU per fly abundances with Acetic and Lactic Acid Bacteria under standard conditions with a combined model.

|  | CFU per Fly |
| --- | --- |
| **Phylum** | **5.0e-59** |
| **Selection Regime** | **6.6e-07** |
| Sex | 0.13 |
| **Phylum x Selection Regime** | **1.5e-28** |
| **Phylum x Sex** | **9.1e-05** |
| Selection Regime x Sex | 0.44 |
| (All) Phylum x Selection x Sex | 0.02 |

**Figure 4:**
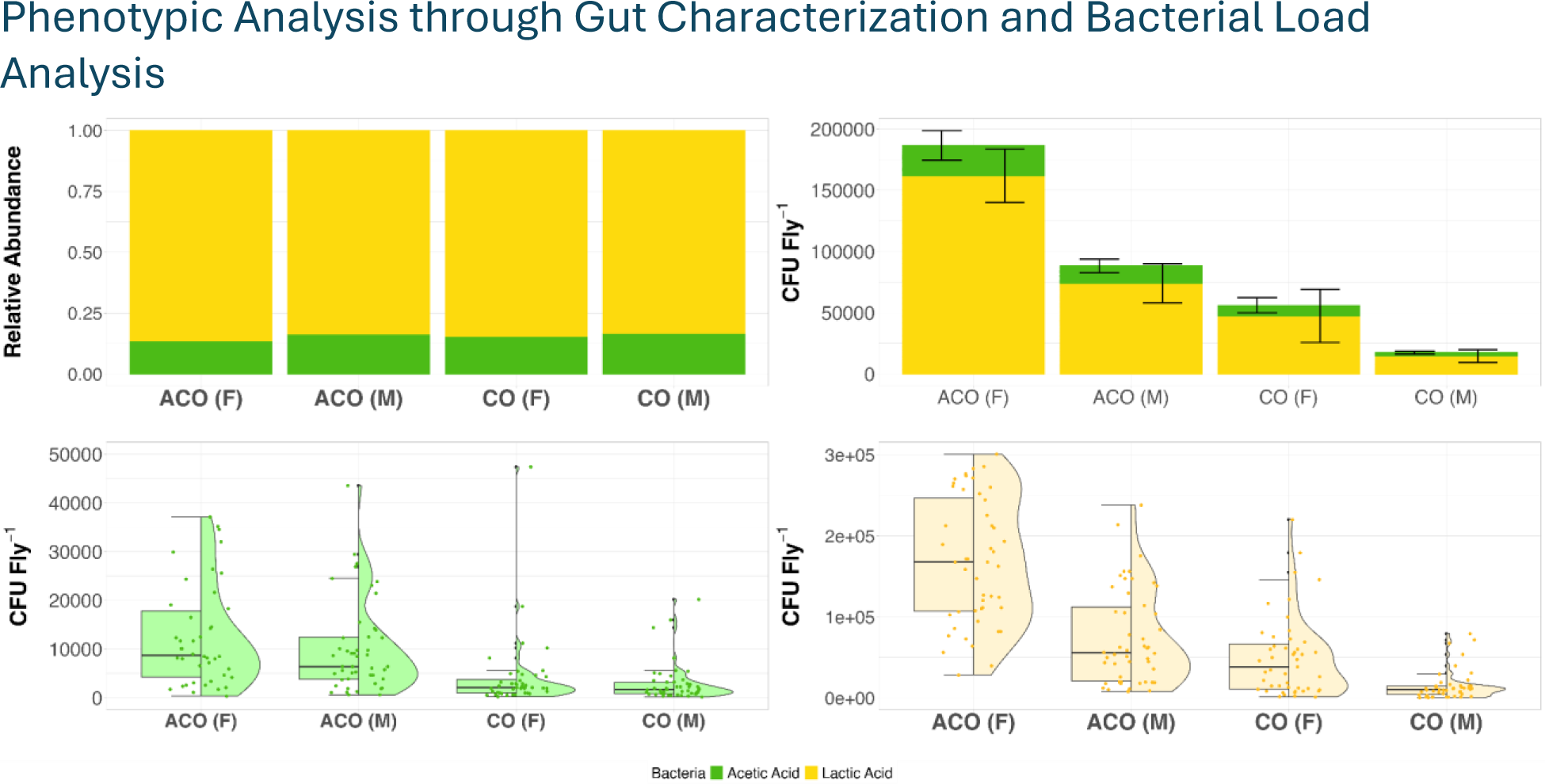
Visualization of Colony Forming Unit (CFU) data under conventional conditions. Top left panel: Relative abundance between acetic acid and lactic acid. Top right panel: absolute CFU abundance per fly. Bottom left panel: Box and violin plot for individual sample abundances of X. Bottom right panel: Box and violin plot for individual sample abundances of X. Note: Four outlier points above 50,000 have been removed for the bottom right panel for graphic fidelity to better capture the density patterns in this visualization. See Supplemental Figure 5 for that graph with all data points included.

Additionally, we investigated the CFU per fly of LAB and AAB in one combined dataset, once again under conditions of raw amounts and amounts normalized by weight (Supplemental Table 2). High significance from the effect of the clade term indicates a selection regime independent and sex independent effect where LAB have a larger abundance than AAB in our samples. We see both a highly significant effect of the interaction term of clade and selection regime as well as a moderately significant effect of selection regime by itself on the levels of bacterial CFUs. ACOs and COs show unique microbiome profiles from one another. While we once again see no effect of sex on CFU per fly by itself, we now see significance from the interaction term of sex and clade, which is something we couldn’t analyze when testing LAB and AAB data separately.

When analyzing the CFU per fly abundances under gnotobiotic conditions (Figure 5), only one term shows only significance across the three models (Supplemental Tables 3); selection regime alone influences the CFU per fly. The Acidic Acid Bacteria model has a p-value of 6.6e-12, the Lactic Acid Bacteria model has a p-value of 0.0014, and the model combining the data into one dataset has a p-value of 5.4e-11. No other terms show significance indicating neither sex nor type of bacteria nor interaction of terms has any impact on the abundance of CFU per fly under gnotobiotic conditions.

**Figure 5:**
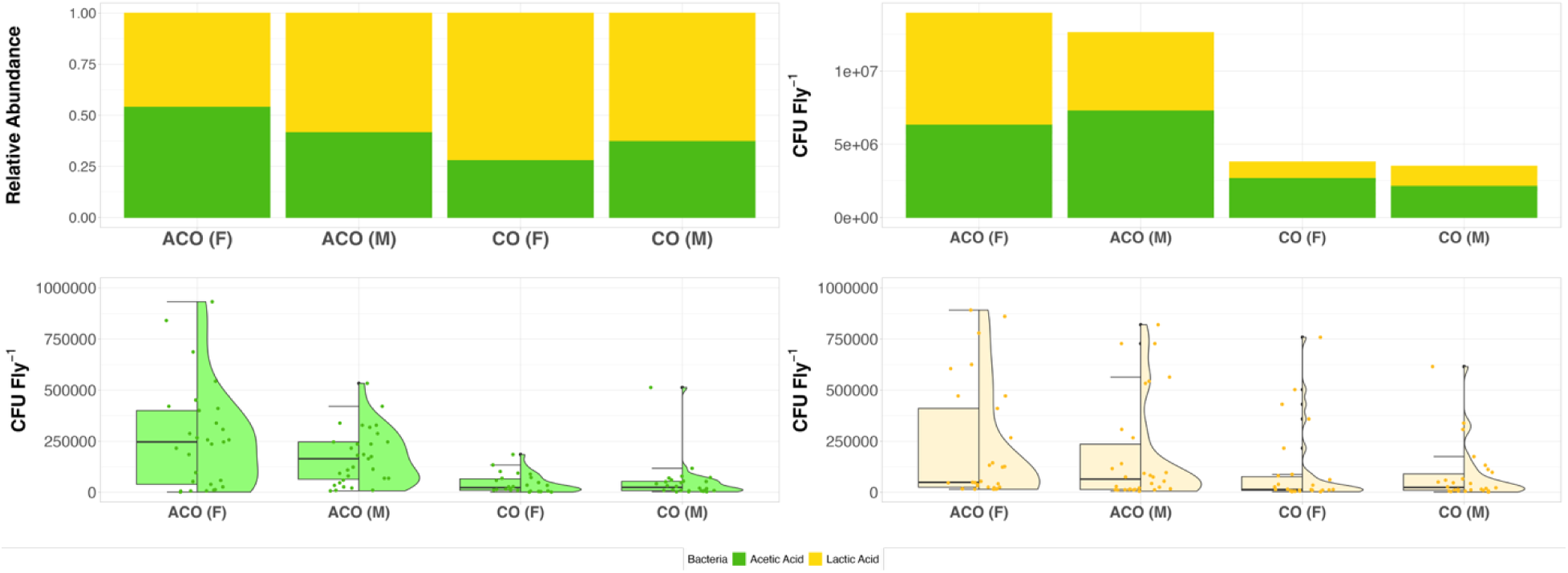
Visualization of Colony Forming Unit (CFU) data under gnotobiotic conditions. Top left panel: Relative abundance between acetic acid and lactic acid. Top right panel: absolute CFU abundance per fly. Bottom left panel: Box and violin plot for population and sex specific abundances of X. Bottom right panel: Box and violin plot for population and sex specific abundances of X. Note: One outlier point above 1,000,000 has been removed for the bottom right panel for graphic fidelity to better capture the density patterns in this visualization.

When using linear models analyzing measurements of gut data between ACO and CO selection regimes (Supplemental Table 4), we find a highly significant different between both the ratio of gut size to body size for the terms of selection regime (*p* = 8.05×10^-49^) as well as sex (*p* = 6.93×10^-27^), but not the interaction term (*p* = 0.158). For the ratio of gut size to abdomen size we find a highly significance effect of selection regime (*p* = 4.53×10^-14^) as well as a modest significant effect of sex (*p* = 2.82×10^-04^), but not the interaction term (*p* = 0.322). Overall, we find that CO flies feature larger body and abdomen sizes then ACO flies. However, ACO flies feature a greater ratio of gut size to body size and gut size to abdomen size. Additionally, female flies have larger body and abdomen sizes than male flies of the same selection regime.

## Discussion

### Metagenomics Analysis of Aging Populations

Our findings provide strong evidence that laboratory selection on life history traits in D. melanogaster leads to stable, repeatable shifts in the composition and abundance of host-associated microbiota. We used 40 populations from the Drosophila Experimental Evolution Population (DEEP) Resource. These populations have undergone laboratory selection for age-of-first reproduction, resulting in significant divergence in life history and physiological traits (Shahrestani et al. 2012, Shahrestani et al. 2016, Burke et al., 2016, Phillips et al., 2018, Kezos et al., 2022). We found that the microbiome profiles in these populations tracked with the evolutionary histories of the populations; populations with closer age-of-first reproduction had more similar microbiomes. When investigating the four most abundant genera in the microbiome, accounting for 99.2% of all bacteria identified across our 40 populations, we observed that although each genotype exhibited a unique microbial signature, genotypes with closer reproductive timing tended to have more similar profiles.

The most pronounced microbiome differences appeared between A-types (populations with the earliest reproduction) and C-types (populations with the latest reproduction). Given that the A- and C-type selection regimes produce nearly a twofold difference in lifespan (Burke et al., 2016), these results suggest that laboratory selection that impacts lifespan is correlated with shifts in the microbiome. In particular, a strong divergence in Wolbachia abundance across populations selected for early (A-type) versus late (C-type) reproduction supports the hypothesis that microbiota composition is not merely a passive outcome of environmental exposure but is additionally under genetic control and itself subject to evolutionary forces.

The most striking and consistent pattern across our datasets was the long-term, stable loss of Wolbachia in A-type populations and its strong persistence in C-types. In both our metagenomic and 16S rRNA sequencing approaches, we found little to no Wolbachia in the A-types and an abundance of Wolbachia in C-types. These findings persisted over hundreds of Drosophila generations, suggesting that host genotype and selection regime shape the colonization and retention of Wolbachia. The clustering of populations by Wolbachia presence closely mirrored divergence in reproductive timing and lifespan, implying that Wolbachia plays a role in, or is influenced by, host life history strategy.

Wolbachia has emerged as a central focus in microbial and evolutionary biology due to its widespread prevalence and profound impacts on host biology. Wolbachia infects an estimated 40% of insect species (Werren et al., 2008), and influences host reproduction, fitness, and pathogen resistance (Bourtzis et al., 1996; Zug & Hammerstein, 2012). In *D. melanogaster*, Wolbachia has been shown to enhance fecundity and provides antiviral protection, likely conferring advantages in slower-developing populations where longevity allows extended reproductive success (Teixeira et al., 2008). While Wolbachia has well-established effects on reproduction and immunity, its functional importance in these populations remains unclear.

Some possibilities emerge to explain the near absence of Wolbachia in A-type populations. First, due to the fast development times of A-type flies, Wolbachia may fail to reach sufficient high densities to be reliably transmitted maternally. Given that Wolbachia transmission is vertical and depends on successful colonization of the female reproductive tract, the accelerated development and early reproduction of A-types may constrain bacterial proliferation before transmission can occur. Second, Wolbachia may confer fitness advantages that become relevant only in genotypes with longer lifespans. For example, Wolbachia has been shown to enhance host immune responses and provide antiviral protection (Kambris et at., 2010, Zhang et al., 2020), benefits that could extend lifespan (Tafesh-Edwards et al., 2024) or improve reproductive output at later ages in C-types, where selection favors the maintenance of high fitness beyond early adulthood. In contrast, the impacts of Wolbachia may be irrelevant - or even costly - in A-types which face no selection for post-reproductive survival and do face selection for early reproduction.

Another plausible explanation for the near absence of Wolbachia in A-type populations is the potential for competitive exclusion by other bacteria taxa, particularly Acetobacter. Prior work has suggested positive, neutral, or negative interactions between members of the microbiota and *Wolbachia* (Imchen et al., 2025, Ye et al., 2017, Simhadri et al., 2017, Fromont et al., 2019, Rota-Stabelli et al., 2023, Fink et al., 2013, Fink et al., 2017). In particular, the acetic acid bacterium *Asaia* has been shown to suppress *Wolbahcia* transmission in mosquitoes (Huges et al., 2014). We do not investigate the mechanisms of transfer here, but high Acetobacter abundance may create an environment that does not support Wolbachia persistence. Supporting this hypothesis is our finding that A-types had higher abundance of Acetobacter compared to C-types. Such competitive dynamics could prevent Wolbachia from establishing or maintaining the population densities necessary for transmission or detection. Some studies have found masking effects during detection of Wolbachia, suggesting that metabolites produced by Acetobacter can interfere with the detection of intracellular Wolbachia, potentially leading to underestimation of Wolbachia abundance in sequencing-based approaches.

Notably, while Wolbachia was undetectable in nearly all A-type populations across multiple generations, we did detect low Wolbachia abundance in single A-type populations from both 2012 and 2018. Interestingly, these were different populations, suggesting that Wolbachia has not been entirely purged from the A-types but instead may persist at cryptic, sub-detectable levels. Importantly, extremely deep-sequencing on a single 2018 A-type population sample maintained zero relative abundance of Wolbachia in accordance with its shallower-read counterpart. These findings point to the potential limitations of standard Illumina-based metagenomic methods for detecting low-abundance intracellular symbionts.

### Phenotypic Analysis through Gut Characterization and Bacterial Load Analysis

Prior research in D. melanogaster suggests that microbial composition can influence gut morphology (Broderick et al., 2014), but there is no clear correlation between gut size, or fly size, with microbial abundance or composition. In the DEEP Resource, fly size is correlated with age-of-first reproduction; populations with early reproduction (A-type_ are smaller than populations with late reproduction (C-type) (Kezos et al., 2023, and in our study). We removed the guts of A-type and C-type flies and measured them.

We found that despite being smaller, A-type flies possess a larger gut-to-body size compared to C-types. This disproportional gut investment may reflect a compensatory mechanism for meeting nutritional and energetic demands under a time-constrained life cycle, but may also relate to microbiome support functions, such as nutrient provisioning, auxotrophy compensation (Conseuga etal., 2020), or microbial contributions to energy metabolism during rapid development (Walters et al., 2020b). Supporting the interpretation that the larger gut-to-body ratios of A-types are not just due to conventional biological necessities, but rather relate to the microbiome is our finding that A-type flies harbor approximately four times as many bacteria per fly as C-types under conventional conditions, despite their smaller body and gut size. This observation aligns with the idea that fast-developing populations may rely heavily on microbial partners to meet immediate survival and reproduction demands.

Results from gnotobiotic experiments reinforce this conclusion. Even when reared under controlled conditions with standardized microbial inocula, A-type flies still hosted significantly higher bacterial loads than C-types. This consistency across environments suggests that differences in bacterial load are driven by host-genotype effects, rather than initial microbiome composition or environment. These findings imply that A-types may either promote microbial proliferation to compensate for limited intrinsic physiological capacity, or alternatively, fail to regulate microbial growth due to reduced investment in immune control or gut homeostasis.

Together, these findings support the growing recognition that the microbiome is not simply a passive passenger but is integrated into the host’s adaptive strategy, potentially contributing to host physiological functions essential for survival and reproduction. A particular compelling direction for future research involves disentangling the temporal sequence of the evolution of host life history traits and the evolution of the host’s ability to genetically control its microbiome. Future studies should investigate whether divergence in life history traits necessitates corresponding shifts in microbiome composition, or whether environmental manipulations (e.g., microbial supplementation) can enhance or buffer the response to laboratory selection on life history traits.

## Conclusion

Our study underscores the critical role of microbiota in shaping and responding to host adaptive evolution. The observed divergence in microbial community composition across *D. melanogaster* populations under different selection-regimes highlights the dynamic interplay between host genetics and microbial ecology. These findings advance our understanding of host-microbiota co-evolution and provide a foundation for exploring and understanding microbial contributions to life history evolution in other systems.

## References

Adair, K.L., Wilson, M., Bost, A., Douglas, A.E., Microbial community assembly in wild populations of the fruit fly *Drosophila melanogaster*, *The ISME Journal*, Volume 12, Issue 4, April 2018, Pages 959–972, 10.1038/s41396-017-0020-x

Bates, D., Mächler, M., Bolker, B., & Walker, S. (2015). Fitting linear mixed-effects models using lme4. Journal of Statistical Software, 67(1), 1–48. 10.18637/jss.v067.i01

Benjamini, Y., & Hochberg, Y. (1995). Controlling the False Discovery Rate: A Practical and Powerful Approach to Multiple Testing. Journal of the Royal Statistical Society: Series B (Methodological), 57(1). 10.1111/j.2517-6161.1995.tb02031.x

Blanco-Míguez, A., Beghini, F., Cumbo, F. et al. Extending and improving metagenomic taxonomic profiling with uncharacterized species using MetaPhlAn 4. Nat Biotechnol 41, 1633–1644 (2023). 10.1038/s41587-023-01688-w

Blum, J. E., Fischer, C. N., Miles, J., & Handelsman, J. (2013). Frequent replenishment sustains the beneficial microbiome of Drosophila melanogaster. mBio, 4(6), e00860–13. 10.1128/mBio.00860-13

Bourtzis, K., Nirgianaki, A., Markakis, G., & Savakis, C. (1996). Wolbachia infection and cytoplasmic incompatibility in Drosophila species. Genetics, 144(3), 1063–1073. 10.1093/genetics/144.3.1063

Bourtzis, K., Pettigrew, M.M. and O’Neill, S.L. (2000), *Wolbachia* neither induces nor suppresses transcripts encoding antimicrobial peptides. Insect Molecular Biology, 9: 635–639. 10.1046/j.1365-2583.2000.00224.x

Broderick, N. A., Buchon, N., & Lemaitre, B. (2014). Microbiota-induced changes in Drosophila melanogaster host gene expression and gut morphology. mBio, 5(3), e01117–14. 10.1128/mBio.01117-14

Burke, M. K., Dunham, J. P., Shahrestani, P., Thornton, K. R., Rose, M. R., & Long, A. D. (2010). Genome-wide analysis of a long-term evolution experiment with Drosophila. Nature, 467(7315), 587–590. 10.1038/nature09352

Burke, M. K., Barter, T. T., Cabral, L. G., Kezos, J. N., Phillips, M. A., Rutledge, G. A., Phung, K. H., Chen, R. H., Nguyen, H. D., Mueller, L. D., & Rose, M. R. (2016). Rapid divergence and convergence of life-history in experimentally evolved Drosophila melanogaster. Evolution; International Journal of Organic Evolution, 70(9). 10.1111/evo.13006

Chandler J.A., Morgan L.J., Bhatnagar S., Eisen J.A., Kopp A. (2011) Bacterial Communities of Diverse Drosophila Species: Ecological Context of a Host–Microbe Model System. PLOS Genetics 7(9): e1002272. 10.1371/journal.pgen.1002272

Chaston J.M., Dobson A.J., Newell P.D., Douglas A.E., (2016). Host Genetic Control of the Microbiota Mediates the Drosophila Nutritional Phenotype. Appl Environ Microbiol 82:. 10.1128/AEM.03301-15

Chippindale, A. K., Chu, T. J. F., & Rose, M. R. (1996). COMPLEX TRADE-OFFS AND THE EVOLUTION OF STARVATION RESISTANCE IN DROSOPHILA MELANOGASTER. Evolution; international journal of organic evolution, 50(2), 753–766. 10.1111/j.1558-5646.1996.tb03885.x

Clark, M. E., Anderson, C. L., Cande, J., & Karr, T. L. (2005). Widespread prevalence of wolbachia in laboratory stocks and the implications for Drosophila research. Genetics, 170(4), 1667–1675. 10.1534/genetics.104.038901

Joseph T. Gale, Rebecca Kreutz, Sarah J. Gottfredson Morgan, Emma K. Davis, Connor Hough, Wendy A. Cisneros Cancino, Brittany Burnside, Ryan Barney, Reese Hunsaker, Ashton Tanner Hoyt, Aubrey Cluff, Maggie Nosker, Chandler Sefcik, Eliza Beales, Jack Beltz, Paul B. Frandsen, Paul Schmidt, John M. Chaston (2024). bioRxiv 2024.10.07.617096; doi: 10.1101/2024.10.07.617096

Fink, C., Staubach, F., Kuenzel, S., Baines, J.F., & Roeder, T. (2013). Noninvasive analysis of microbiome dynamics in the fruit fly Drosophila melanogaster. Applied and Environmental Microbiology, 79(22):6984–8. doi: 10.1128/AEM.01903-13.

Fink, C., Von Frieling, J., Knop, M., & Roeder, T. (2017). Drosophila fecal sampling. Bio-Protocol, 7(18):e2547. doi: 10.21769/BioProtoc.2547

Fromont, C., Adair, K.L., & Douglas, A.E. (2019). Correlation and causation between the microbiome, Wolbachia and host functional traits in natural populations of drosophilid flies. Molecular Ecology, 28(7):1826–1841. doi: 10.1111/mec.15041

Graves, J. L., Jr, Hertweck, K. L., Phillips, M. A., Han, M. V., Cabral, L. G., Barter, T. T., Greer, L. F., Burke, M. K., Mueller, L. D., & Rose, M. R. (2017). Genomics of Parallel Experimental Evolution in Drosophila. Molecular biology and evolution, 34(4), 831– 842. 10.1093/molbev/msw282

Hughes, G. L., Dodson, B. L., Johnson, R. M., Murdock, C. C., Tsujimoto, H., Suzuki, Y., Patt, A. A., Cui, L., Nossa, C. W., Barry, R. M., Sakamoto, J. M., Hornett, E. A., & Rasgon, J. L. (2014). Native microbiome impedes vertical transmission of Wolbachia in Anopheles mosquitoes. Proceedings of the National Academy of Sciences of the United States of America, 111(34), 12498–12503. 10.1073/pnas.1408888111

Imchen, M., Kaur, R., Zhou Luo, D.H., Bitman, M.W., Lefoulon, E., McGarry, A., Bordenstein, S.R., & Bordenstein, S.R. (2024). Gut-Germline Axis: A Reproductive Endosymbiont’s Adaptation is Modulated by the Gut Microbiome. bioRxiv. doi: 10.1101/2024.09.10.611936

Ives P. T. (1970). FURTHER GENETIC STUDIES OF THE SOUTH AMHERST POPULATION OF DROSOPHILA MELANOGASTER. Evolution; international journal of organic evolution, 24(3), 507–518. 10.1111/j.1558-5646.1970.tb01785.x

Kassambara, A. (2023). rstatix: Pipe-Friendly Framework for Basic Statistical Tests. R package version 0.7.2, https://rpkgs.datanovia.com/rstatix/.

Kezos, J. N., Barter, T. T., Phillips, M. A., Cabral, L. G., Greenspan, Z. S., Arnold, K. R., Azatian, G., Buenrostro, J., Bhangoo, P. S., Khong, A., Reyes, G. T., Rahman, A., Humphrey, L. A., Bradley, T. J., Mueller, L. D., & Rose, M. R. (2023). Building Bridges from Genome to Physiology Using Machine Learning and Drosophila Experimental Evolution. Physiological and Biochemical Zoology, 96(3). 10.1086/724827

Kambris, Z., Blagborough, A. M., Pinto, S. B., Blagrove, M. S., Godfray, H. C., Sinden, R. E., & Sinkins, S. P. (2010). Wolbachia stimulates immune gene expression and inhibits plasmodium development in Anopheles gambiae. PLoS pathogens, 6(10), e1001143. 10.1371/journal.ppat.1001143

Kolde, R. (2022). Package “pheatmap”: Pretty heatmaps. R Package

Kozich, J. J., Westcott, S. L., Baxter, N. T., Highlander, S. K., & Schloss, P. D. (2013). Development of a dual-index sequencing strategy and curation pipeline for analyzing amplicon sequence data on the MiSeq Illumina sequencing platform. Applied and Environmental Microbiology, 79(17), 5112–5120. 10.1128/AEM.01043-13

Phillips, M. A., Rutledge, G. A., Kezos, J. N., Greenspan, Z. S., Talbott, A., Matty, S., Arain, H., Mueller, L. D., Rose, M. R., & Shahrestani, P. (2018). Effects of evolutionary history on genome wide and phenotypic convergence in Drosophila populations. BMC Genomics, 19,743. 10.1186/s12864-018-5118-7

Rose, M. R., & Charlesworth, B. (1980). A test of evolutionary theories of senescence. Nature, 287(5772), 141–142. 10.1038/287141a0

Rose, M. R. (1984). Laboratory evolution of postponed senescence in Drosophila melanogaster. Evolution, 38(5), 1004–1010. 10.2307/2408434

Team, R. C. (2023). R Core Team 2023 R: A language and environment for statistical computing. R foundation for statistical computing. https://www.R-project.org/. *R Foundation for Statistical Computing*.

Rota-Stabelli, O., Houle, D., & Müller, H. (2023). Wolbachia endosymbionts alter the gut microbiome in Drosophila nigrosparsa. Environmental Microbiology, 25(5):1867– 1880. doi: 10.1111/1462-2920.16271 (PMID: 36958642)

Rudman, S.M., Greenblum, S., Hughes, R.C., Rajpurohit, S., Kiratli, O., Lowder, D.B., Lemmon, S.G., Petrov, D.A., Chaston, J.M., & Schmidt, P. (2019). Microbiome composition shapes rapid genomic adaptation of *Drosophila melanogaster*, Proc. Natl. Acad. Sci. U.S.A. 116 (40) 20025–20032, 10.1073/pnas.1907787116.

Simhadri, R.K., Fast, E.M., Guo, R., Schultz, M.J., Vaisman, N., Ortiz, L., Bybee, J., Slatko, B.E., & Frydman, H.M. (2017). The gut commensal microbiome of Drosophila melanogaster is modified by the endosymbiont Wolbachia. mSphere, 2:e00287–17. doi: 10.1128/mSphere.00287-17

Staubach F, Baines JF, Künzel S, Bik EM, Petrov DA (2013) Host Species and Environmental Effects on Bacterial Communities Associated with Drosophila in the Laboratory and in the Natural Environment. PLOS ONE 8(8): e70749. 10.1371/journal.pone.0070749

Tafesh-Edwards, G., Kyza Karavioti, M., Markollari, K., Bunnell, D., Chtarbanova, S., & Eleftherianos, I. (2024). Wolbachia endosymbionts in Drosophila regulate the resistance to Zika virus infection in a sex dependent manner. Frontiers in microbiology, 15, 1380647. 10.3389/fmicb.2024.1380647

Teixeira, L., Ferreira, Á., & Ashburner, M. (2008). The bacterial symbiont Wolbachia induces resistance to RNA viral infections in Drosophila melanogaster. PLoS Biology, 6(12), e1000002. 10.1371/journal.pbio.1000002

Werren, J. H., Baldo, L., & Clark, M. E. (2008). Wolbachia: Master manipulators of invertebrate biology. Nature Reviews Microbiology, 6(10), 741–751. 10.1038/nrmicro1969

Wickham, H. (2016). ggplot2 Elegant Graphics for Data Analysis (Use R!). Springer.

Wong, C.N.A., Ng, P. and Douglas, A.E. (2011), Low-diversity bacterial community in the gut of the fruitfly Drosophila melanogaster. Environmental Microbiology, 13: 1889–1900. 10.1111/j.1462-2920.2011.02511.x

Wong, C.N.A., Chaston, J.M., Douglas, A.E., (2013) The inconstant gut microbiota of *Drosophila* species revealed by 16S rRNA gene analysis, *The ISME Journal*, Volume 7, Issue 10, Pages 1922–1932, 10.1038/ismej.2013.86

Ye, Y.H., Seleznev, A., Flores, H.A., Woolfit, M., & McGraw, E.A. (2017). Gut microbiota in Drosophila melanogaster interacts with Wolbachia but does not contribute to Wolbachia-mediated antiviral protection. Journal of Invertebrate Pathology, 143:18–25. doi: 10.1016/j.jip.2016.11.011

Yu Y, Iatsenko I. (2025). Drosophila symbionts in infection: when a friend becomes an enemy. Infect Immun 93:e00511–24. 10.1128/iai.00511-24 https://doi.org/10.1128/iai.00511-24

Zhang, D., Wang, Y., He, K., Yang, Q., Gong, M., Ji, M., & Chen, L. (2020). Wolbachia limits pathogen infections through induction of host innate immune responses. PloS one, 15(2), e0226736. 10.1371/journal.pone.0226736

Zug, R., & Hammerstein, P. (2012). Still a host of hosts for Wolbachia: Analysis of recent data suggests that 40% of terrestrial arthropod species are infected. PLoS One, 7(6), e38544. 10.1371/journal.pone.0038544

